# Urban Establishment of *Aedes vittatus* in Yucatan, Mexico

**DOI:** 10.1101/2025.10.17.678774

**Authors:** Juan Navarrete-Carballo, Carlos Arisqueta-Chablé, Wilbert Bibiano-Marín, Jorge Palacio-Vargas, Marvin Lugo-Moguel, Jimmy Torres-Castro, Herón Huerta, Azael Che-Mendoza, Fabián Correa-Morales, Gonzalo M Vazquez-Prokopec, Judith Ortega-Canto, Pablo Manrique-Saide

**Author notes:** Address for correspondence: Pablo Manrique-Saide, Universidad Autónoma de Yucatán, Unidad Colaborativa para Bioensayos Entomológicos, Campus de Ciencias Biológicas y Agropecuarias, Km. 15.5 Carretera Mérida-Xmatkuiil, CP. 97315, Mérida, México.

## Abstract

*Aedes vittatus*, an invasive mosquito competent for multiple arboviruses, has expanded far beyond its historical range. We report its first detection and urban establishment in mainland Americas, in Merida, Mexico. This underscores the urgent need to strengthen surveillance, risk assessment, and preparedness against invasive *Aedes* species across the region.

## Text

*Aedes*-borne arboviruses—including dengue, Zika and chikungunya, remain persistent public health challenges across the Americas. While *Aedes aegypti* is the primary vector and *Ae. albopictus* a secondary one, the emergence of additional invasive species presents new challenges to sustainable vector control. *Aedes vittatus*, widely distributed across Africa and Asia, is implicated in yellow fever transmission and recognized as competent for dengue, Zika and chikungunya based on experimental and field studies (*1,2*). Over the past decade, its range expanded into southern Europe (Spain, France, Italy, Portugal) (*2,3*). In the Americas, detections had been limited to Caribbean islands (Dominican Republic, Cuba and Jamaica) (*2,4–6*), but its arrival in continental areas raises major concerns for arbovirus surveillance and control.

We report the local establishment of *Ae. vittatus* in Merida, Mexico, representing the first confirmed record of this invasive species established in a mainland urban area of the Americas. During routine malaria-vector surveillance by the Yucatan Ministry of Health, a blood-fed female *Ae. vittatus* was collected at Laguna San Marcos (20°54′57″N, 89°39′54″W) during evening human landing catches (HLC) on July 22, 2025. Following this finding, intensive surveys were conducted (July–August 2025) in three adjacent neighborhoods: San Marcos Nocoh I, Roble Agrícola, and Gran Roble (Fig. 1). Collections included late-afternoon HLC, BG-Sentinel, and CDC light traps, along with inspections of immature habitats. A transect of 30 houses near the pond was surveyed, including Prokopack aspirations indoors and outdoors.

**Figure 1.**
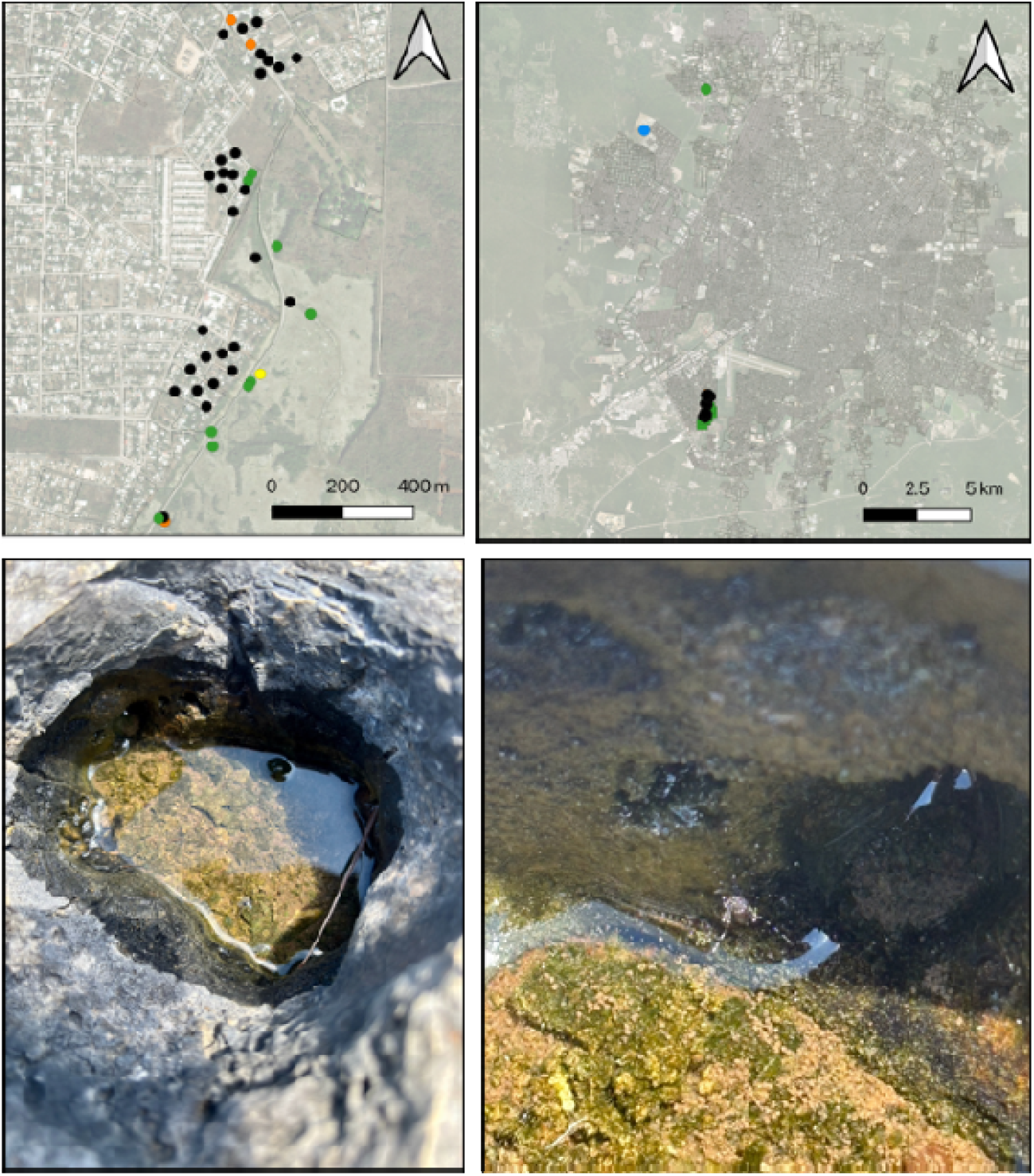
A) Location of survey and collection sites for *Aedes vittatus* in Merida, Yucatan (blue points = positive for larvae; green = positive for adults; yellow = positive for larvae and pupae; orange = positive for larvae and adults; black = negative for *Ae. vittatus*). B) Location of additional recent collection sites of *Ae. vittatus* in the metropolitan area of Merida. C) Natural rock pool temporarily filled with rainwater. D) Adult *Aedes vittatus* emerging from a natural breeding site in Merida, Yucatan.

Across collections, 1,428 mosquitoes representing 15 species from 5 genera were identified. Among them, 38 *Ae. vittatus* adults (27♀ [3 fed, 24 unfed], 11♂) were captured by HLC (n=15), BG-Sentinel (n=2), CDC light traps (n=15), and Prokopacks (n=6). Additionally, 35 immatures (30 larvae, 5 pupae) were recovered from rock pools, pond margins, a rain-filled boat, and animal drinking containers. *Ae. vittatus* accounted for 5.1% of all specimens, compared with *Ae. aegypti* (31.1%) and *Ae. albopictus* (1.7%). Of 30 houses inspected, 26 were positive for adult mosquitoes. *Ae. vittatus* adults (n=6; 5♀ unfed, 1♂) were found only outdoors in peridomestic areas of 4 premises. Of 45 containers surveyed, 12 harbored immatures. In households, 20 larvae were recovered (18 from an animal drinking container, 2 from a small boat). At the nearby pond, 32 adults and 15 immatures were also collected. Species identifications using taxonomic keys (*7,8*) were validated by the National Laboratory at InDRE, where voucher specimens were deposited (Fig. 2).

**Figure 2.**
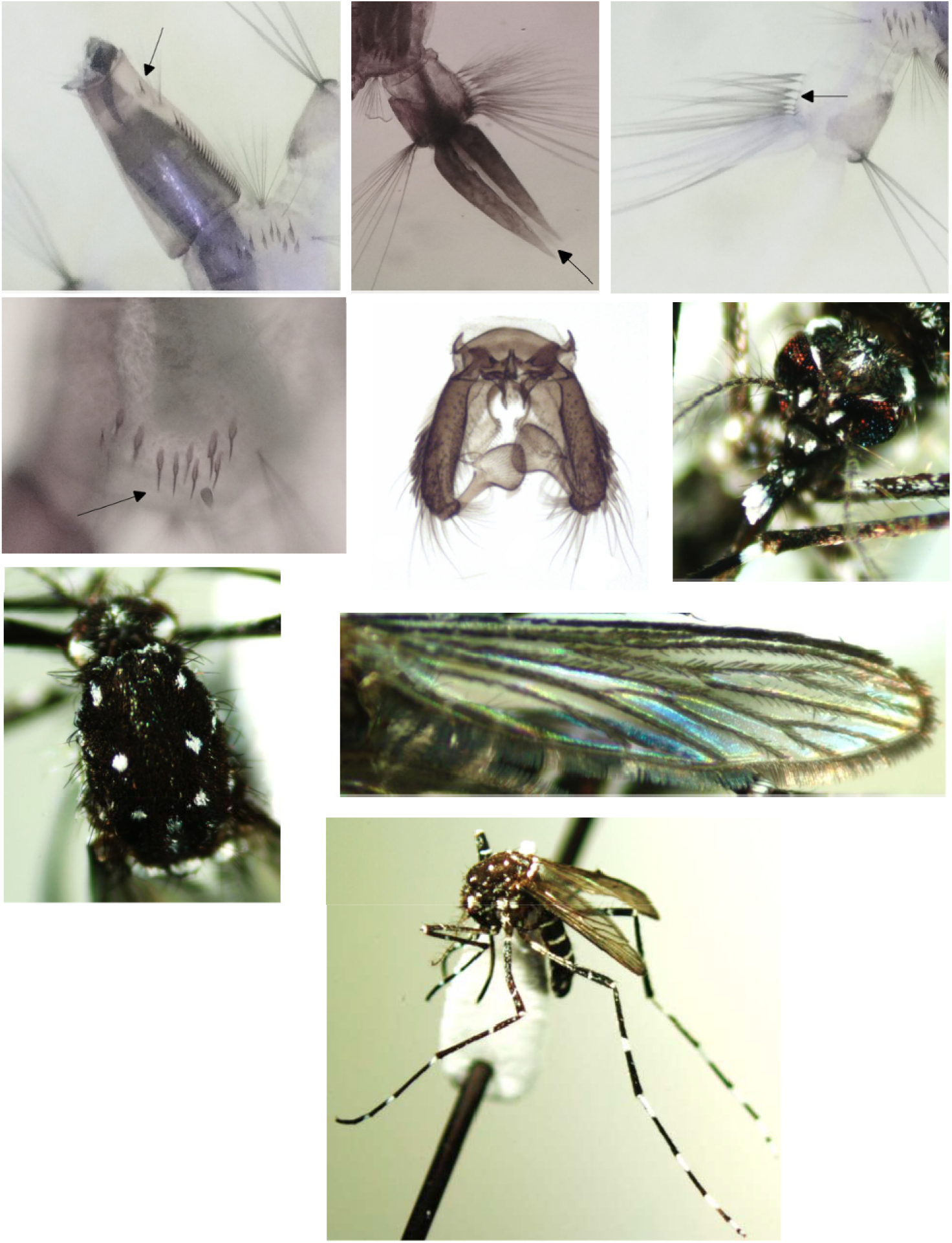
Diagnostic characters of *Aedes vittatus*. Larval characters. A) Siphon (acus absent) with a pair of isolated pecten teeth located just beyond the middle of the siphon, posterior to seta 1-S. B) Anal gills pointed, longer than the saddle. C) Segment X with precratal hairs. D) Comb of abdominal segment VIII arranged in an irregular row with 9–12 teeth. Adults. E) Male *Ae. vittatus*, genitalia in ventral view. F) Female *Ae. vittatus*, head in frontal view. G) Thorax in dorsal view. H) Wing in lateral view, I) Female *Ae. vittatus* pinned.

Detection of larvae, pupae, and adults of both sexes across diverse habitats indicates local establishment rather than transient introduction (*9,10*). Recovery of multiple developmental stages within a 1.5 km radius demonstrates a self-sustaining population. Within 4–6 weeks, larvae were also detected 13.8 km away, and an adult female was collected 15.4 km from the original site, confirming dispersal within the metropolitan area. Upon confirmation, the Yucatan Ministry of Health alerted national authorities (CENAPRECE), which issued a national entomologic surveillance alert. Locally, ovitrap monitoring and intensified adult collections were implemented, along with immediate control measures such as handheld thermal fogging and vehicle-mounted space spraying.

In its native range, *Ae. vittatus* breeds mainly in rock and tree holes, but during its expansion it has colonized artificial containers, showing high ecological plasticity. Our findings confirm this adaptability, with collections from both natural and artificial habitats. Given Merida’s abundance of container habitats, including rock holes, conditions are favorable for peri-urban and urban establishment. Possible ecological interactions with *Ae. aegypti*, the dominant local species, warrant study. Mexico’s national ovitrapping network (209,096 traps in 396 municipalities) provides a robust platform for early detection of invasive container-breeding *Aedes*. In Merida, 5,998 ovitraps are active and are expected to help clarify *Ae. vittatus* distribution within Yucatan and beyond in the near future.

In 2023, Yucatan reported >10,000 dengue cases, the highest incidence in Mexico, with Merida accounting for ≈45%. The coexistence of *Ae. aegypti, Ae. albopictus*, and now *Ae. vittatus* raises concerns about shifts in arbovirus transmission dynamics. The capture of blood-fed females during HLC suggests some degree of anthropophily, consistent with previous reports. However, no adults were collected indoors, suggesting absence of endophily. The future role of *Ae. vittatus* in urban arbovirus transmission in Mexico and the Americas remains poorly defined. Nevertheless, its establishment in a tropical urban area endemic for various arboviruses highlights the need to incorporate this species into systematic surveillance, including entomologic monitoring and xenomonitoring, to assess transmission potential. Early detection of invasive *Aedes* species has proven critical in Europe and the Caribbean. The establishment of *Ae. vittatus* in Merida underscores the importance of strengthening national surveillance networks and fostering regional collaboration across the Americas.

## Disclaimers

The authors declare no conflicts of interest. The authors used ChatGPT to improve clarity and suggest title and running head options. All content was subsequently reviewed and edited by the authors, who take full responsibility for the final publication.

## Author Bio

Juan Navarrete-Carballo is an entomologist at the Yucatan Ministry of Health, assigned to the Collaborative Unit for Entomological Bioassays at the Universidad Autónoma de Yucatán. His research focuses on the biology and taxonomy of arthropod vectors of public health importance.

## Notes

### Competing Interest Statement

The authors have declared no competing interest.

